# Genetic structure of the stingless bee *Tetragonisca angustula*

**DOI:** 10.1101/026740

**Authors:** Flávio O. Francisco, Leandro R. Santiago, Yuri M. Mizusawa, Benjamin P. Oldroyd, Maria C. Arias

**Author notes:** Correspondence: Flávio O. Francisco, Departamento de Genética e Biologia Evolutiva, Instituto de Biociências, Universidade de São Paulo, Rua do Matão 277 – sala 320, 05508-090, São Paulo, SP, Brazil.

## Abstract

The stingless bee *Tetragonisca angustula* Latreille 1811 is distributed from Mexico to Argentina and is one of the most widespread bee species in the Neotropics. However, this wide distribution contrasts with the short distance traveled by females to build new nests. Here we evaluate the genetic structure of several populations of *T. angustula* using mitochondrial DNA and microsatellites. These markers can help us to detect differences in the migratory behavior of males and females. Our results show that the populations are highly differentiated suggesting that both females and males have low dispersal distance. Therefore, its continental distribution probably consists of several cryptic species.

## Introduction

The stingless bee *Tetragonisca angustula* Latreille 1811 distributed from Mexico to Argentina is one of the most widespread bee species in the Neotropics (Silveira *et al.*, 2002; Camargo & Pedro, 2013). It is a small (4-5 mm in length), generalist and highly eusocial bee (Michener, 2007) and highly adaptable to different nest sites. Colonies comprise up to 5,000 individuals (Lindauer & Kerr, 1960), and are usually built in tree trunks or in wall cavities. It swarms frequently and is extremely successful in urban environments (Batista *et al.*, 2003; Slaa, 2006; Velez-Ruiz *et al.*, 2013). *Tetragonisca angustula* is one of the most popular stingless bees for meliponiculture in Latin America (Nogueira-Neto, 1997; Cortopassi-Laurino *et al.*, 2006) and nest transportation and trading is very common among beekeepers.

In general, colony reproduction in stingless bees begins by workers searching for a new nest site within their foraging range (van Veen & Sommeijer, 2000a). Daughter nests are established at most a few hundred meters from the “mother” nest (Nogueira-Neto, 1997). After selecting the site, several workers begin to transport cerumen, propolis and honey from the mother nest to the new one (Nogueira-Neto, 1997). This nest site preparation phase can last from a few days (van Veen & Sommeijer, 2000a) to a few months (Nogueira-Neto, 1997). A virgin queen then leaves the mother nest accompanied by hundreds of workers (van Veen & Sommeijer, 2000b). The next day the virgin queen flies out, mates with presumably one male (Peters *et al.*, 1999; Palmer *et al.*, 2002), returns to the nest and about a week later begins oviposition (van Veen & Sommeijer, 2000b).

In contrast, little is known about the reproductive behavior of stingless bee males. After emergence from brood cells, they remain in the nest for two to three weeks (Cortopassi-Laurino, 2007). They then leave the nest and never return. There are no data about the behavior of males during their period outside the nest. In the laboratory, males can live up to six weeks (Velthuis *et al.*, 2005). Therefore, they likely have two to four weeks for dispersal and reproduction. Males often form mating aggregations, and these are comprised of males from hundreds of different, and not necessarily nearby colonies (Paxton, 2000; Cameron *et al.*, 2004; Kraus *et al.*, 2008; Mueller *et al.*, 2012). This suggests high male dispersal.

Studies on the genetic structure of populations can help us better understand dispersal behavior and evolutionary history. There are three existing population genetic studies of *T. angustula* (Oliveira *et al.*, 2004; Baitala *et al.*, 2006; Stuchi *et al.*, 2008). These studies are of limited scope due to limited sampling or the use of outdated molecular markers such as RAPDs and isozymes. Thus, the conclusions are ambiguous concerning the extent and direction of gene flow, population differentiation, dispersal and evolutionary history.

Here we evaluate the genetic structure of several populations of *T. angustula* using mitochondrial DNA (mtDNA) and microsatellite markers. Considering the wide distribution of *T. angustula* and the commonness of nest transportation and trading, we expect low genetic differentiation among populations despite the low dispersal distance of females during swarming.

## Materials and methods

### Sampling

We collected 1,002 *T. angustula* from 457 sites distributed on the mainland and islands in south/south-eastern Brazil (Table S1). Eleven islands all with arboreal vegetation and of area greater than 1.0 km^2^ were selected, 10 being land-bridge islands isolated about 12,000 years ago (Suguio *et al.*, 2005) and one sedimentary island (Ilha Comprida) which arose about 5,000 years ago (Suguio *et al.*, 2003). The islands range in size from 1.1 to 451 km^2^ and are 0.1 to 38 km from the mainland (Table S2, Fig. 1). Bees were sampled from nests (*n* = 125, one per nest) and flowers (*n* = 877) (Table S1). Samples were grouped into 17 populations, 14 from the mainland and three from islands (Fig. 1).

**Fig. 1.**
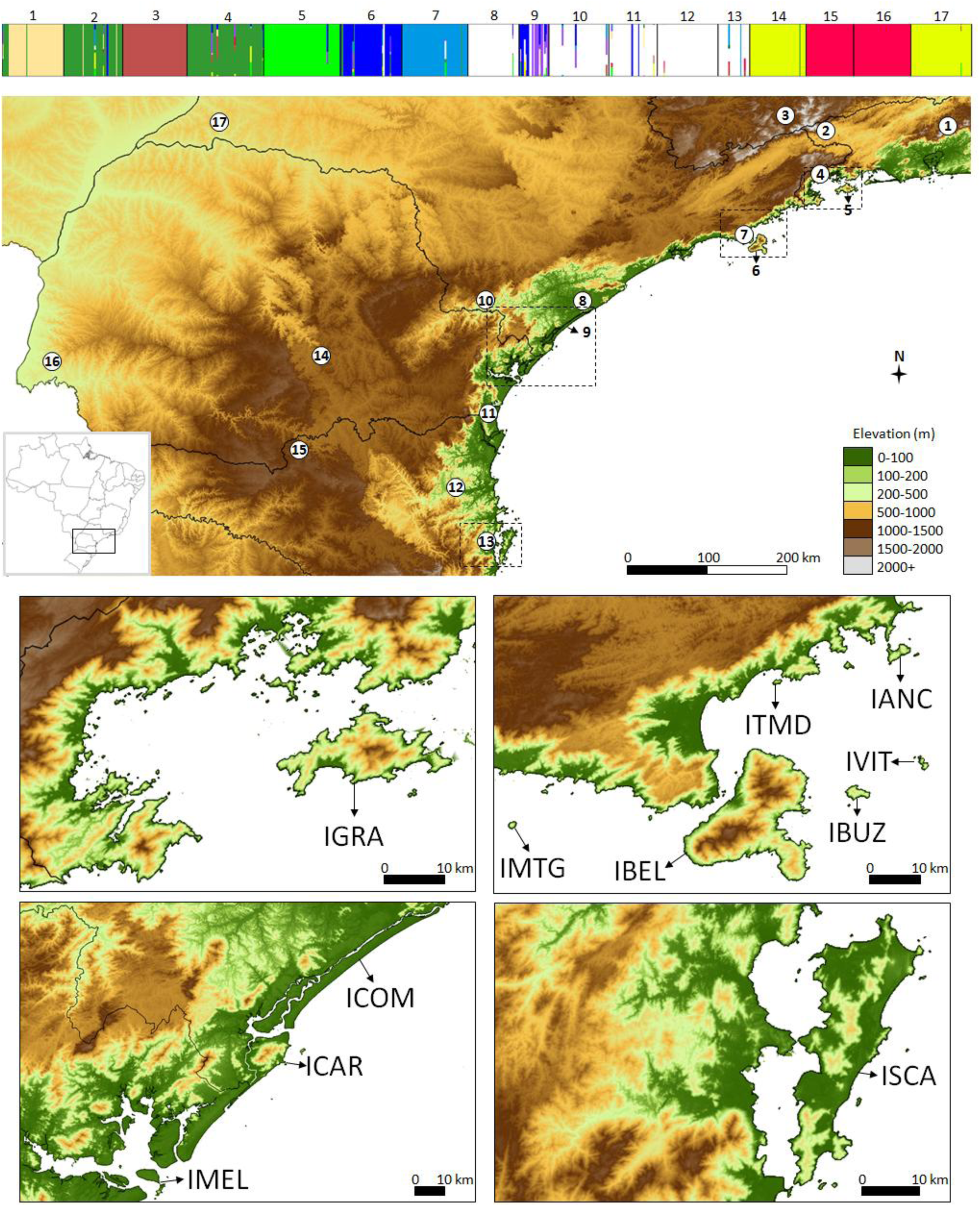
Posterior probability assignment (vertical axis) of individual genotypes (horizontal axis) for *K* = 10 (*Tetragonisca angustula*) according to the program BAPS (upper panel). Below, map of the studied area with the approximate location of the sampled populations. Population names are 1: Teresópolis, 2: Resende, 3: Passa Quatro, 4: Angra dos Reis, 5: Ilha Grande, 6: São Sebastião, 7: Ilha de São Sebastião, 8: Iguape, 9: Ilha Comprida, 10: Apiaí, 11: Guaratuba, 12: Blumenau, 13: São José, 14: Prudentópolis, 15: Porto União, 16: Foz do Iguaçu, and 17: Teodoro Sampaio. Detailed location of all islands visited (lower panels). IGRA: Ilha Grande; IANC: Ilha Anchieta; ITMD: Ilha do Tamanduá; IVIT: Ilha da Vitória; IBUZ: Ilha de Búzios; IBEL: Ilha de São Sebastião; IMTG: Ilha Monte de Trigo. ICOM: Ilha Comprida; ICAR: Ilha do Cardoso; IMEL: Ilha do Mel. ISCA: Ilha de Santa Catarina.

We preserved the specimens in 96% ethanol for transport to the laboratory. DNA extraction followed the protocol described in Francisco *et al.* (2014). We dried the specimens at room temperature for 20 min prior to DNA extraction. 83

### Mitochondrial DNA sequencing

Two mitochondrial genes were partially sequenced: cytochrome c oxidase subunit 1 (*COI*) and cytochrome b (*Cytb*). Details about amplification and sequencing are given in Francisco *et al.* (2014).

### Microsatellite genotyping

The samples were genotyped for eleven microsatellite loci: Tang03, Tang11, Tang12, Tang17, Tang29, Tang57, Tang60, Tang65, Tang68, Tang70, and Tang77 (Brito *et al.*, 2009). PCR conditions for each locus are given in Francisco *et al.* (2014). Electrophoresis, visualization and genotyping were performed according to Francisco *et al.* (2011).

Micro-checker 2.2.3 (van Oosterhout *et al.*, 2004) was used to identify null alleles and scoring errors. Colony 2.0.1.7 (Jones & Wang, 2010) was used to determine whether individuals collected in the same plant or places nearby were related. Samples were excluded from our data set if matched all of the following three criteria: collected at sites distant less than 2 km, indicated as related by colony, and sharing a mtDNA haplotype. Overall, 722 *T. angustula* bees from 17 populations were deemed suitable for further genetic analyses (Table 1).

**Table 1.**
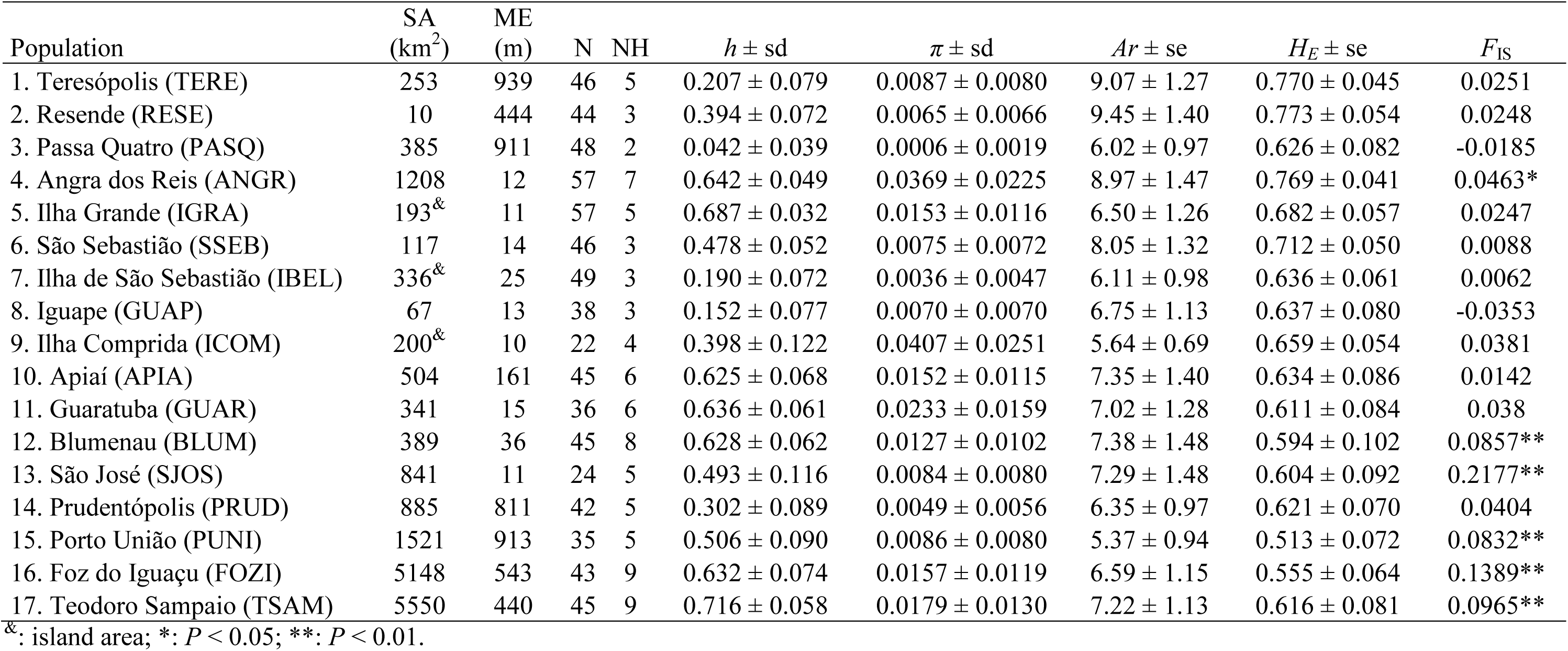
Population characteristics and genetic diversity in *Tetragonisca angustula* populations. SA: sampled area in square kilometers. ME: median elevation in meters. N: sample size. NH: number of haplotypes. *h* ± sd: haplotype diversity and standard deviation. *π* ± sd: nucleotide diversity and standard deviation. *Ar* ± se: allelic richness after rarefaction for 22 individuals and standard error. *H_E_* ± se: expected heterozigosity and standard error. *F*_IS_: inbreeding coefficient.

| Population | SA<br>(km<sup>2</sup>) | ME<br>(m) | N | NH | $h \pm sd$ | $\pi \pm sd$ | $Ar \pm se$ | $H_E \pm se$ | $F_{IS}$ |
| --- | --- | --- | --- | --- | --- | --- | --- | --- | --- |
| 1. Teresópolis (TERE) | 253 | 939 | 46 | 5 | $0.207 \pm 0.079$ | $0.0087 \pm 0.0080$ | $9.07 \pm 1.27$ | $0.770 \pm 0.045$ | 0.0251 |
| 2. Resende (RESE) | 10 | 444 | 44 | 3 | $0.394 \pm 0.072$ | $0.0065 \pm 0.0066$ | $9.45 \pm 1.40$ | $0.773 \pm 0.054$ | 0.0248 |
| 3. Passa Quatro (PASQ) | 385 | 911 | 48 | 2 | $0.042 \pm 0.039$ | $0.0006 \pm 0.0019$ | $6.02 \pm 0.97$ | $0.626 \pm 0.082$ | -0.0185 |
| 4. Angra dos Reis (ANGR) | 1208 | 12 | 57 | 7 | $0.642 \pm 0.049$ | $0.0369 \pm 0.0225$ | $8.97 \pm 1.47$ | $0.769 \pm 0.041$ | 0.0463* |
| 5. Ilha Grande (IGRA) | 193<sup>&</sup> | 11 | 57 | 5 | $0.687 \pm 0.032$ | $0.0153 \pm 0.0116$ | $6.50 \pm 1.26$ | $0.682 \pm 0.057$ | 0.0247 |
| 6. São Sebastião (SSEB) | 117 | 14 | 46 | 3 | $0.478 \pm 0.052$ | $0.0075 \pm 0.0072$ | $8.05 \pm 1.32$ | $0.712 \pm 0.050$ | 0.0088 |
| 7. Ilha de São Sebastião (IBEL) | 336<sup>&</sup> | 25 | 49 | 3 | $0.190 \pm 0.072$ | $0.0036 \pm 0.0047$ | $6.11 \pm 0.98$ | $0.636 \pm 0.061$ | 0.0062 |
| 8. Iguape (GUAP) | 67 | 13 | 38 | 3 | $0.152 \pm 0.077$ | $0.0070 \pm 0.0070$ | $6.75 \pm 1.13$ | $0.637 \pm 0.080$ | -0.0353 |
| 9. Ilha Comprida (ICOM) | 200<sup>&</sup> | 10 | 22 | 4 | $0.398 \pm 0.122$ | $0.0407 \pm 0.0251$ | $5.64 \pm 0.69$ | $0.659 \pm 0.054$ | 0.0381 |
| 10. Apiaí (APIA) | 504 | 161 | 45 | 6 | $0.625 \pm 0.068$ | $0.0152 \pm 0.0115$ | $7.35 \pm 1.40$ | $0.634 \pm 0.086$ | 0.0142 |
| 11. Guaratuba (GUAR) | 341 | 15 | 36 | 6 | $0.636 \pm 0.061$ | $0.0233 \pm 0.0159$ | $7.02 \pm 1.28$ | $0.611 \pm 0.084$ | 0.038 |
| 12. Blumenau (BLUM) | 389 | 36 | 45 | 8 | $0.628 \pm 0.062$ | $0.0127 \pm 0.0102$ | $7.38 \pm 1.48$ | $0.594 \pm 0.102$ | 0.0857** |
| 13. São José (SJOS) | 841 | 11 | 24 | 5 | $0.493 \pm 0.116$ | $0.0084 \pm 0.0080$ | $7.29 \pm 1.48$ | $0.604 \pm 0.092$ | 0.2177** |
| 14. Prudentópolis (PRUD) | 885 | 811 | 42 | 5 | $0.302 \pm 0.089$ | $0.0049 \pm 0.0056$ | $6.35 \pm 0.97$ | $0.621 \pm 0.070$ | 0.0404 |
| 15. Porto União (PUNI) | 1521 | 913 | 35 | 5 | $0.506 \pm 0.090$ | $0.0086 \pm 0.0080$ | $5.37 \pm 0.94$ | $0.513 \pm 0.072$ | 0.0832** |
| 16. Foz do Iguaçu (FOZI) | 5148 | 543 | 43 | 9 | $0.632 \pm 0.074$ | $0.0157 \pm 0.0119$ | $6.59 \pm 1.15$ | $0.555 \pm 0.064$ | 0.1389** |
| 17. Teodoro Sampaio (TSAM) | 5550 | 440 | 45 | 9 | $0.716 \pm 0.058$ | $0.0179 \pm 0.0130$ | $7.22 \pm 1.13$ | $0.616 \pm 0.081$ | 0.0965** |

<sup>&</sup>: island area; \*: $P < 0.05$ ; \*\*: $P < 0.01$ .

Genepop 4.1.2 (Rousset, 2008) was used to verify Hardy-Weinberg equilibrium (HWE) in populations and loci and to detect linkage disequilibrium (LD). Markov chain was set for 10,000 dememorizations, 1,000 batches and 10,000 iterations per batch. In cases of multiple comparisons, *P*-values were corrected by applying Sequential Goodness of Fit test by the program Sgof 7.2 (Carvajal-Rodríguez *et al.*, 2009). This method is advantageous over other correction methods because it increases its statistical power with the increasing of the number of tests (Carvajal-Rodríguez *et al.*, 2009).

### Genetic diversity

Arlequin 3.5.1.3 (Excoffier & Lischer, 2010) was used to calculate mtDNA haplotype (*h*) and nucleotide (*π*) diversity. Genalex 6.5 (Peakall & Smouse, 2006, 2012) was used to calculate microsatellite allelic richness (*A*) and expected heterozygosity (*HE*). Since sample sizes were different, allelic richness was standardized by rarefaction (*Ar*) using the program Hp-rare 1.0 (Kalinowski, 2005). Differences in *Ar* among populations were estimated by Mann-Whitney two-tailed U Test (Mann & Whitney, 1947). Inbreeding coefficients (*F*_IS_) were calculated for each population with 10,000 permutations using Arlequin.

### Population differentiation and gene flow

Mega 5.2.1 (Tamura *et al.*, 2011) was used to calculate the number of base substitutions per site from mitochondrial sequences by averaging over all sequence pairs between populations using the Kimura 2-parameter (K2p) model (Kimura, 1980). Population pairwise *θ* values (an *F*_ST_ analogue, Weir & Cockerham 1984) were calculated with 10,000 permutations by arlequin using microsatellite alleles. When heterozygosity is high, *F*_ST_ and its analogues may not be appropriate measures of genetic differentiation (Hedrick, 2005; Jost, 2008; Heller & Siegismund, 2009). For this reason, Jost’s *D*_est_ (Jost, 2008) was calculated. This statistc is not influenced by heterozygosity (Jost, 2008) and is more appropriate for microsatellite data (Heller & Siegismund, 2009). Global *D*_est_ was calculated with 9,999 permutations for mtDNA and microsatellite data using Genalex. Pairwise *D*_est_ was calculated only for microsatellite data. Mantel tests between genetic and geographical distances among populations were performed with 9999 permutations by Genalex to verify isolation by distance for both molecular markers.

Baps 6 (Corander *et al.*, 2008; Cheng *et al.*, 2013) was used to infer population structure using microsatellites and the geographic coordinates of the sampled individuals to spatially cluster them. Baps 6 provides a Bayesian analysis of genetic population structure that creates *K* groups of individuals based on the similarity of their genotypes. The program was initially ran 5 times for each of *K* = 1 to 17 and then 10 times for each of *K* = 5 to 14. These results were used for admixture analysis with 200 iterations to estimate the admixture coefficients for the individuals, 200 simulated reference individuals per population and 20 iterations to estimate the admixture coefficients of the reference individuals.

Estimates of rates and direction of current and/or recent migration (*m*) between populations were determined by the program bayesass 3 (Wilson & Rannala, 2003) using microsatellites multilocus genotypes through Markov chain Monte Carlo (MCMC) techniques. We performed five independent runs with 10^7^ MCMC iterations, burn-in of 10^6^ iterations and sampling frequency of 2,000. The delta values used were 0.25 (migration), 0.40 (allele frequencies) and 0.55 (inbreeding).

### Assessment of population demography

To detect any recent bottleneck events we used the program bottleneck 1.2.02 (Piry *et al.*, 1999). We used the two-phased model (TPM) of mutation which is suggested as the most appropriate for microsatellites (Di Rienzo *et al.*, 1994). The variance among multiple steps was 12 and the proportion of stepwise mutation model in the TPM was 95% as suggested by Piry *et al.* (1999). Altogether 10,000 iterations were performed. The significance of any deviation was determined with a Wilcoxon sign-rank test.

## Results

### Island occurrences

*Tetragonisca angustula* was found and collected on five of the 11 islands visited (Table 1). However, only the samples from Ilha Grande, Ilha de São Sebastião and Ilha Comprida were included in the analyses. The other collections were not included due to small sample size (Ilha do Cardoso, *n* = 1), and to individuals being highly related, with anecdotal reports of introduced nests (Ilha de Santa Catarina, see Francisco *et al.* (2014)).

### MtDNA diversity 164

The *COI* gene sequences were 417 bp long (GenBank accession numbers KF222891-KF223893) and 32 haplotypes were identified. The *Cytb* sequences were 391 bp long (KF223894-KF224896) and generated 43 haplotypes. Most differences among haplotypes were synonymous substitutions, since the number of distinct amino acid sequences were four for *COI* and 15 for *Cytb*. We concatenated the nucleotide sequences (808 bp) for population analyses.

The 722 concatenated sequences defined 73 haplotypes. Since *h* and *π* were positively correlated (*r* = 0.510, *P* = 0.036, *n* = 17) we hereafter use *π* as our measure of mtDNA diversity. Nucleotide diversity ranged from 0.0006 ± 0.0019 (Passa Quatro) to 0.0407 ± 0.0251 (Ilha Comprida) (Table 1).

There was a non-significant positive correlation between the size of the sampled area and mtDNA diversity (*r* = 0.135, *P* = 0.606, *n* = 17). The correlation between median elevation and mtDNA diversity was negative but non-significant (*r* = −0.428, *P* = 0.087, *n* = 17).

### MtDNA differentiation

Population structure was high. Of 73 haplotypes 67 were population-specific. We built a haplotype network where the frequency and distribution of haplotypes are shown (Fig. S1). The network shows a ‘star-pattern’ centered on four haplotypes. It illustrates the high number of endemic haplotypes, and the great number of nucleotide substitutions that separate the Porto União/Foz do Iguaçu populations from the others. The populations of Teresópolis, Resende, Prudentópolis, Angra dos Reis, and Ilha Grande all feature unique haplotypes.

Global *D*_est_ was 0.772 (*P* < 0.001) indicating a highly significant population structure. The highest K2p values were found for Porto União/Foz do Iguaçu with respect to all other populations (2.809% to 3.306%) (Table 2).

**Table 2.**
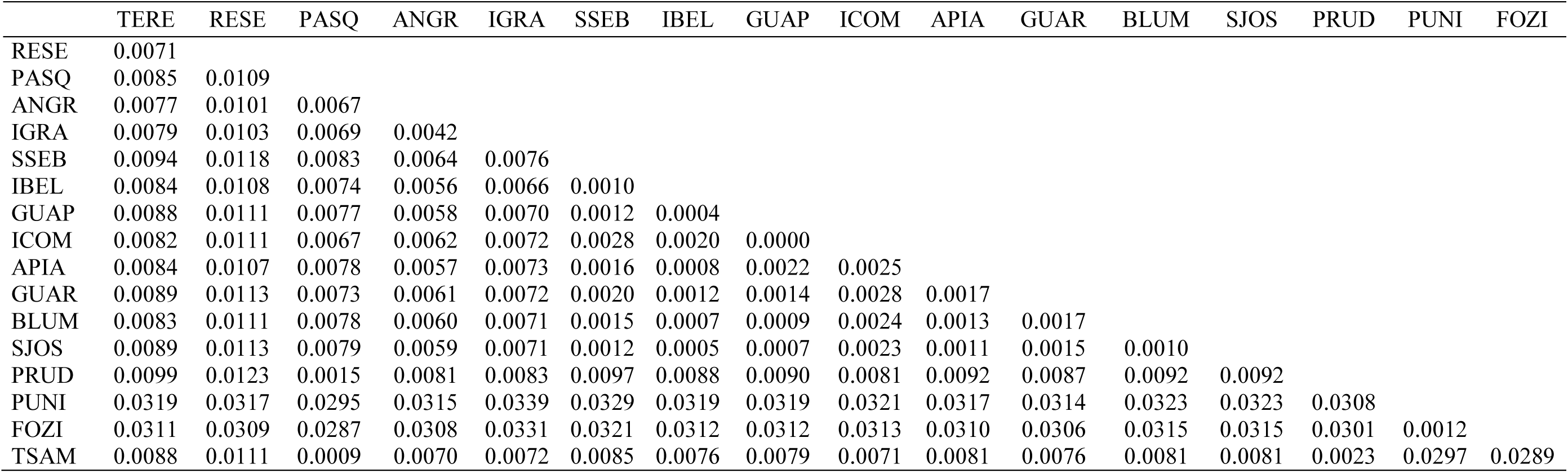
Estimates of evolutionary divergence over sequence pairs between populations of *Tetragonisca angustula*. The number of base substitutions per site from averaging over all sequence pairs between populations are shown. Analyses were conducted using the Kimura 2-parameter model (Kimura, 1980) and involved 722 nucleotide sequences. Population abbreviations as in Table 1.

|  | TERE | RESE | PASQ | ANGR | IGRA | SSEB | IBEL | GUAP | ICOM | APIA | GUAR | BLUM | SJOS | PRUD | PUNI | FOZI |
| --- | --- | --- | --- | --- | --- | --- | --- | --- | --- | --- | --- | --- | --- | --- | --- | --- |
| RESE | 0.0071 |  |  |  |  |  |  |  |  |  |  |  |  |  |  |  |
| PASQ | 0.0085 | 0.0109 |  |  |  |  |  |  |  |  |  |  |  |  |  |  |
| ANGR | 0.0077 | 0.0101 | 0.0067 |  |  |  |  |  |  |  |  |  |  |  |  |  |
| IGRA | 0.0079 | 0.0103 | 0.0069 | 0.0042 |  |  |  |  |  |  |  |  |  |  |  |  |
| SSEB | 0.0094 | 0.0118 | 0.0083 | 0.0064 | 0.0076 |  |  |  |  |  |  |  |  |  |  |  |
| IBEL | 0.0084 | 0.0108 | 0.0074 | 0.0056 | 0.0066 | 0.0010 |  |  |  |  |  |  |  |  |  |  |
| GUAP | 0.0088 | 0.0111 | 0.0077 | 0.0058 | 0.0070 | 0.0012 | 0.0004 |  |  |  |  |  |  |  |  |  |
| ICOM | 0.0082 | 0.0111 | 0.0067 | 0.0062 | 0.0072 | 0.0028 | 0.0020 | 0.0000 |  |  |  |  |  |  |  |  |
| APIA | 0.0084 | 0.0107 | 0.0078 | 0.0057 | 0.0073 | 0.0016 | 0.0008 | 0.0022 | 0.0025 |  |  |  |  |  |  |  |
| GUAR | 0.0089 | 0.0113 | 0.0073 | 0.0061 | 0.0072 | 0.0020 | 0.0012 | 0.0014 | 0.0028 | 0.0017 |  |  |  |  |  |  |
| BLUM | 0.0083 | 0.0111 | 0.0078 | 0.0060 | 0.0071 | 0.0015 | 0.0007 | 0.0009 | 0.0024 | 0.0013 | 0.0017 |  |  |  |  |  |
| SJOS | 0.0089 | 0.0113 | 0.0079 | 0.0059 | 0.0071 | 0.0012 | 0.0005 | 0.0007 | 0.0023 | 0.0011 | 0.0015 | 0.0010 |  |  |  |  |
| PRUD | 0.0099 | 0.0123 | 0.0015 | 0.0081 | 0.0083 | 0.0097 | 0.0088 | 0.0090 | 0.0081 | 0.0092 | 0.0087 | 0.0092 | 0.0092 |  |  |  |
| PUNI | 0.0319 | 0.0317 | 0.0295 | 0.0315 | 0.0339 | 0.0329 | 0.0319 | 0.0319 | 0.0321 | 0.0317 | 0.0314 | 0.0323 | 0.0323 | 0.0308 |  |  |
| FOZI | 0.0311 | 0.0309 | 0.0287 | 0.0308 | 0.0331 | 0.0321 | 0.0312 | 0.0312 | 0.0313 | 0.0310 | 0.0306 | 0.0315 | 0.0315 | 0.0301 | 0.0012 |  |
| TSAM | 0.0088 | 0.0111 | 0.0009 | 0.0070 | 0.0072 | 0.0085 | 0.0076 | 0.0079 | 0.0071 | 0.0081 | 0.0076 | 0.0081 | 0.0081 | 0.0023 | 0.0297 | 0.0289 |

### Microsatellite diversity

After the Sequential Goodness of Fit correction, deviation from HWE was occasional, likely arising from type 1 error, and therefore no locus was removed from the analyses (Table S3). No significant LD was found between any pair of loci (all *P* > 0.05).

Microsatellite diversity was moderate to high. *Ar* and *H_E_* were positively correlated (*r* = 0.787, *P* < 0.001, *n* = 17). Hereafter we use *Ar* as our measure of microsatellite diversity. *Ar* was standardized for 22 individuals and ranged from 5.37 (Porto União) to 9.45 (Resende) (Table 1). *Ar* was significantly different only between Porto União and Resende (*U* = 93, *P* = 0.033) and Porto União and Teresópolis (*U* = 29, *P* = 0.039).

There was a negative but non-significant correlation between *Ar* and size of the sampled area (*r* = -0.114, *P* = 0.662, *n* = 17) and between *Ar* and median elevation (*r* = −0.084, *P* = 0.748, *n* = 17).

Six populations had inbreeding coefficients (*F*_IS_) significantly different from zero (*P* < 0.05). The highest *F*_IS_ (0.2177) was found in São José (Table 1).

### Microsatellite differentiation

Global *D*_est_ was high (0.375, *P* < 0.001) and indicates population structure. Pairwise comparisons also detected population structure, since most *θ* values were between 0.05 and 0.15 (Table S4) and most *D*_est_ values were higher than 0.25 (Table 3). Pairwise *θ* and *D*_est_ were positively correlated (*r* = 0.977, *P* < 0.001, *n* = 136) and we use *D*_est_ as our measure of microsatellite differentiation hereafter. *D*_est_ ranged from 0.0204 (Guaratuba × Blumenau) to 0.8464 (Prudentópolis × Foz do Iguaçu). High *D*_est_ values were always detected in comparisons between Porto União/Foz do Iguaçu and other populations. Low differentiation was observed in some populations near the coast (Iguape, Apiaí, Guaratuba, Blumenau, and São José) but also inland (Porto União × Foz do Iguaçu and Prudentópolis × Teodoro Sampaio).

**Table 3.**
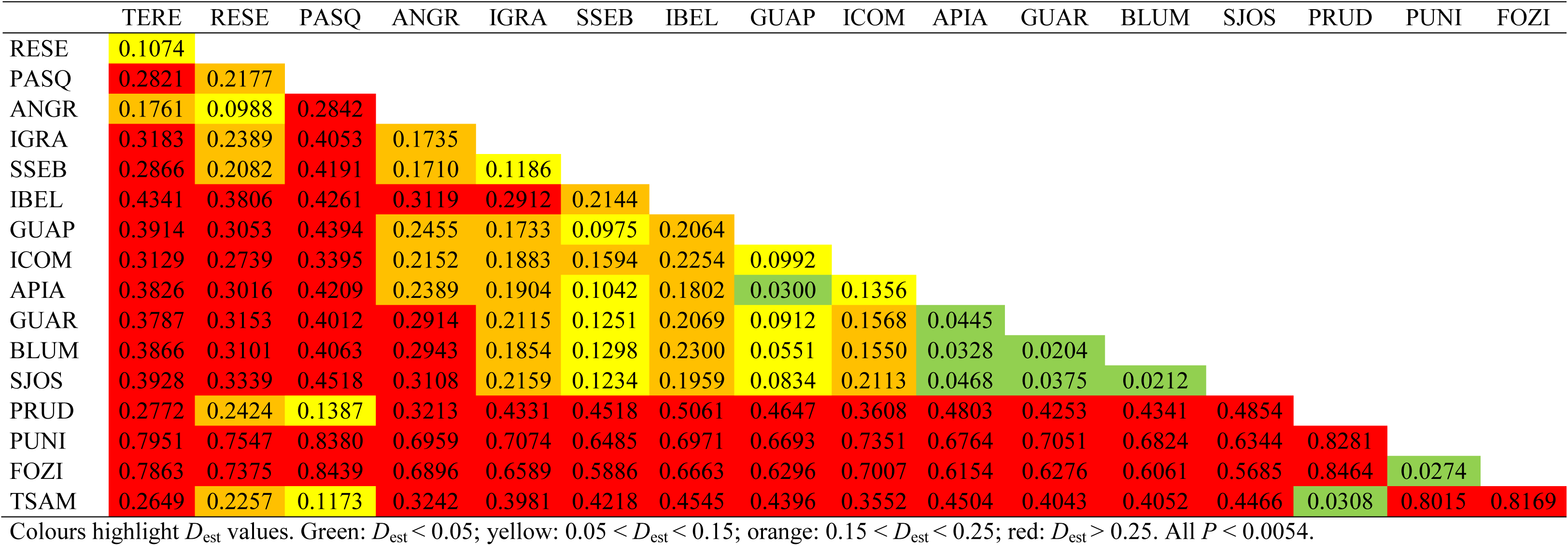
Pairwise index of differentiation (*D*_est_) from microsatellite data of *Tetragonisca angustula*. Population abbreviations as in Table 1.

Population structure was also suggested by the spatial cluster approach used by BAPS, which determined *K* = 10 as the most likely optimal number of clusters (probability of 98.99%). The clusters were [Foz do Iguaçu/Porto União], [Iguape/Apiaí/Guaratuba/ Blumenau/SãoJosé], [Ilha Comprida], [São Sebastião], [Ilhabela], [Ilha Grande], [Passa Quatro], [Teodoro Sampaio/Prudentópolis], [Teresópolis], [Resende/Angra dos Reis] (Fig. 1). *D*_est_ results are in good agreement with these clusters.

The results of the migration rates estimated in bayesass suggested a low level of gene flow throughout the studied area (Table S5). Only 15 out of 272 comparisons showed *m* > 0 between two populations. Most of the populations that showed evidence of gene flow are near the coast (Fig. 1), but inland populations such as Porto União × Foz do Iguaçu also showed evidence of gene flow. Migration is directional. For instance, the non-differentiation detected between Prudentópolis × Teodoro Sampaio is apparently due to a high migration rate (0.2486 migrants per generation) from Prudentópolis to Teodoro Sampaio, whereas migration in the opposite direction was not detected (Table S5). The results obtained by bayesass are in good agreement with the population structure indicated by *θ*, *D*_est_ and baps.

### Isolation by distance

There was a positive and significant correlation between geographic and genetic distance for both mitochondrial (*r* = 0.415, *P* = 0.004, *n* = 136) and microsatellite markers (*r* = 0.464, *P* < 0.001, *n* = 136).

### Population demography

We did not detect recent bottlenecks in any of the 17 populations (all *P* > 0.1392, Table S6).

## Discussion

Our results show that *T. angustula* populations are highly differentiated as demonstrated by mtDNA and microsatellite markers. This suggests that both females and males have low dispersal distance.

Per population mtDNA nucleotide diversity (*π*) ranged from low to high. High *π* suggests that a population had a long evolutionary history and a large effective population size. Low *π* may be explained by lineage sorting or suggest that a population bottleneck has occurred in the past (Avise, 2000). The characteristic star shape of the *T. angustula* haplotype network provides evidence of relatively recent local extinction, re-colonization, and population expansion. Several phylogeographic studies of vertebrate and invertebrate populations, conducted in some of the areas that we studied, also found low mtDNA nucleotide diversity (Cabanne *et al.*, 2007; Carnaval *et al.*, 2009; Batalha-Filho *et al.*, 2010; Brito & Arias, 2010; Francisco & Arias, 2010; D’Horta *et al.*, 2011; Bell *et al.*, 2012). As argued in these papers, changes in sea level during the Pleistocene generated population bottlenecks followed by species expansion, and this is reflected in localized low nucleotide diversity to this day. Therefore, it is likely that populations that have high mtDNA diversity (e.g. Angra dos Reis) did not experience recent bottlenecks, while populations with low mtDNA diversity (e.g. Passa Quatro) are in regions that likely arose by a recent population expansion.

Overall, we found high mitochondrial genetic differentiation between populations. Similar population structuring has been observed for other stingless bee species (Brito & Arias, 2010; Francisco & Arias, 2010; Quezada-Euán *et al.*, 2012; Brito *et al.*, 2013; Francisco *et al.*, 2013). The mtDNA population structure of stingless bees probably arises from their reproductive behavior. Nonetheless, some populations are not well differentiated from others. This is likely due to gene natural flow, although human transportation also likely plays a role. For instance, haplotypes 34, 35, and 36 all found in Ilha Comprida, were similar to those found in Passa Quatro/Teodoro Sampaio (34 and 36) and Teresópolis (35) (Fig. S1). Due to the high frequency of endemic haplotypes, and the physical distance between these populations, we suggest that nests have been transported to Ilha Comprida causing an artificial increase in this population’s mtDNA diversity.

Nuclear genetic diversity was moderate to high in all populations. Microsatellite diversity was not significantly different between populations except for Porto União. This result shows that the ecological features of each sampling site are not influencing the molecular diversity. Indeed, variables such as size of the sampled area and median elevation were not significantly correlated with genetic diversity for both mtDNA and microsatellites. Moreover, our results did not detect recent bottleneck (e.g. due to habitat fragmentation) in any of the studied populations. However, it is worth emphasizing that theoretical and practical studies have shown that habitat fragmentation affects immediately the genetic structure by increasing it, while the reduction of genetic diversity may take longer (Varvio *et al.*, 1986; Keyghobadi *et al.*, 2005). According to this statement, it will be necessary to monitor the genetic diversity of *T. angustula* populations studied over years, since currently high structure was detected.

Evidence of inbreeding was found in six populations (Angra dos Reis, Blumenau, São José, Porto União, Foz do Iguaçu and Teodoro Sampaio). This might be an artifact caused by Wahlund effect (Hartl & Clark, 2007) and/or a consequence of the low dispersal of *T. angustula*. If the latter is true, the persistence of these populations is at risk (Keller & Waller, 2002).

Microsatellite data also indicated high genetic structuring and low gene flow among populations. This suggests that like females, the dispersal distance of males is also quite limited even between populations separated by 34 km of continuous forest. It is interesting here to note that all island populations were differentiated from their mainland counterparts, indicating that males do not cross water for distances as short as 300 m. The program BAYESASS suggested that the highest migration rate is from Prudentópolis to Teodoro Sampaio, populations separated by more than 300 km. These two populations do not share any mtDNA haplotypes suggesting that this gene flow is mediated only by males following the stepping stone model (Kimura & Weiss, 1964).

For both markers population clusters appear to be unrelated to physical barriers (such as rivers or mountain ranges) or forest presence, indicating that genetic connectivity demands more than just habitat connectivity (Marsden *et al.*, 2012). Populations may diverge even when there are no apparent obstacles to gene flow due to low dispersal, geographic distance and genetic drift (isolation by distance). Overall, the population structure of *T. angustula* is shaped by isolation by distance.

The highest genetic divergence observed was between Porto União/Foz do Iguaçu and the remaining populations. At least 15 mtDNA mutation steps separate these two populations from the others. This represents about 2.8 to 3.3% divergence, which is as high as the divergence between lineages A and Y of *Apis mellifera* (Franck *et al.*, 2001), which are thought to have diverged over one million years ago (Whitfield *et al.*, 2006). Francisco *et al.* (2014) suggested that bees from Porto União and Foz do Iguaçu might belong to the subspecies *T. angustula fiebrigi* while the others to *T. angustula angustula.*

Among the islands we visited only Ilha do Mel (Zanella, 2005), Ilha de Santa Catarina (Steiner *et al.*, 2006) and Ilha Grande (Lorenzon *et al.*, 2006) had been previously surveyed for bees and *T. angustula* was reported on all of them. We did not locate *T. angustula* on six of the 11 islands we visited. Our failure to verify *T. angustula* on most islands may be due its ancestral absence on the islands when they became isolated or to its extinction after isolation. The constraint on queen dispersal prevents (re)colonization of islands whose distance from the mainland is greater than a few hundred of meters. Even if (re)colonization has occurred, its establishment may not have been successful. With low dispersal, *T. angustula* has low effective population size and high extinction rate. Island size may be critical to the survival of viable *T. angustula* populations – we were unable to locate them on any island less than 28 km_2_. Competition among colonies doubtless limits the number of colonies an island can support so that small islands may not be able to maintain viable populations of *T. angustula*. The rarity of stingless bee species on islands has been noted elsewhere (Schwartz-Filho & Laroca, 1999; Zanella, 2005).

Our results indicate that *T. angustula* is not genetically homogeneous across the studied area. Considering that this species has a continental distribution, we speculate this species is ancient and includes wide range of genetically different taxa with the same (or similar) morphology. Sampling across its entire distribution range is needed to elucidate its taxonomic status as well as its evolutionary history.

## Acknowledgments

We are grateful to Paulo Henrique P. Gonçalves for his help with the sampling and to Susy Coelho and Julie Lim for technical assistance. We thank Adílson de Godoy, Carlos Chociai, Flávio Haupenthal, Geraldo Moretto, Marcos Wasilewski, Marcos Antonio, Renato Marques, José Moisés, André Trindade, Teófilo, Eduardo da Silva, Guaraci Cordeiro, Marcos Fujimoto, PC Fernandes, Samuel Boff, Thaiomara Alves, the managers and the staff of the Parks, the residents of Ilha da Vitória, Ilha de Búzios and Ilha Monte de Trigo, and countless people who assisted us in the fieldwork. We thank Dr. Jeffrey Lozier for comments on an early version of this manuscript. For permits, we thank Instituto Brasileiro do Meio Ambiente e dos Recursos Naturais Renováveis (IBAMA) and Instituto Chico Mendes de Conservação da Biodiversidade (ICMBio) (18457-1), Instituto Florestal do estado de São Paulo (260108 - 000.000.002.517/0 2008), Instituto Ambiental do estado do Paraná (128/09) and Instituto Estadual do Ambiente do Rio de Janeiro (E-07/300.011/0). This work was supported by Fundação de Amparo à Pesquisa do Estado de São Paulo (04/15801-0; 08/07417-6; 08/08546-4; 10/18716-4; 10/50597-5) and Australian Research Council. This work was developed in the Research Center on Biodiversity and Computing (BioComp) of the Universidade de São Paulo (USP), supported by the USP Provost’s Office for Research.

